# Transforming off-the-shelf personal glucose meter into a sustainable and decentralized label-free nucleic acid and NAAT detection platform

**DOI:** 10.64898/2026.05.16.725651

**Authors:** Aditya Chourasia, Saba Parveen, Shrawan Kumar, Arunansu Talukdar, Mrittika Sengupta, Souradyuti Ghosh

**Affiliations:** Centre for Life Sciences, Mahindra University, Hyderabad, Telangana, India; Interdisciplinary Centre for Nanosensors and Nanomedicine, Mahindra University, Hyderabad, Telangana, India; Department of Geriatric Medicine, Medical College and Hospital Kolkata, Kolkata

**Keywords:** Nucleic acid sensor, NAAT detection, Label-free, Glucometer, Point-of-care device

## Abstract

In today’s world, point-of-care nucleic acid detection still remains extensively constrained and limited by the heavy dependence on centralized urban instrumentation facilities and complex assay workflows. Here, we elucidate a glucometer-based analytical platform that enables label-free detection of nucleic acids and the nucleic acid amplification products through a simple redox-mediated mechanism. The approach leverages the potassium ferricyanide (K_3_[Fe(CN)_6_])/ potassium ferrocyanide (K_4_[Fe(CN)_6_]), redox system, which is intrinsic to commercial glucometers, complementing with interactions between methylene blue (MB) and nucleic acids. These interactions transduce concentration differences in nucleic acids into quantifiable electrochemical signal readouts. Distinct varied signal outputs are observed between single-stranded and double-stranded DNA, enabling the direct detection as well as integration with nucleic acid amplification tests (NAATs), including polymerase chain reaction, rolling circle amplification, and loop-mediated isothermal amplification. Optimization of reaction parameters and conditions leads to enhancement of the overall signal discrimination and sensitivity across various assay formats. This innovation repurposes widely available off-the-shelf glucometers as a low-cost, portable nucleic acid detectors, thus eliminating the need for any specialized instrumentation. Our results enumerate and establish a generalized and scalable strategy for nucleic acid sensing. The platform thus supports sustainable and environmentally responsible point-of-care testing, thereby enabling improved accessibility and public health monitoring at resource-limited and remote settings.

## Introduction

Point-of-care (PoC) nucleic acid testing is an increasingly crucial and evolving for rapid diagnostics, especially in resource-limited settings and far-off remote areas. Nucleic acid detection via NAAT allows sensitive detection of pathogenic diseases or biomarkers for diseases easily and inexpensively. On the other hand, conventional methods like PCR require inadequately expensive, bulky instruments as well as highly skilled and trained operators. However, meeting the WHO’s REASSURED criteria which is-Real-time connectivity, Ease of specimen collection, Affordable, Sensitive, Specific, User-friendly, Rapid and robust, Equipment-free or simple, and Deliverable to end-users is critical for PoC diagnostics. Among biosensors, electrochemical PoC sensors fulfill several of the REASSURED criteria: they use simple and less complex equipment, respond efficiently and rapidly, and yield a high sensitivity and specificity even in complex and higher order samples^1^. Unlike lab-based assays, electrochemical DNA/NAAT platforms can be highly miniaturized and portable; they operate on very small sample volumes with immediate and cost-effective readouts^1^. Nevertheless, typical electrochemical setups are still dependent on custom electrodes and potentiostats, which can be highly expensive and technical proficiency demanding. To overcome these shortcomings, one promising strategy involves harnessing off-the-shelf glucometers that have underwent significant technological refinements to become increasingly accurate, user-friendly, cost-effective, and portable analytical devices^2^. Commercial glucometers are globally distributed, calibrated and regulated, and intrinsically low-cost (INR 500 – 1500, roughly USD 5–15 per unit). Using them as generic biosensors has potential leverages for mass manufacturing and miniaturization to reduce costs and improve sustainability and circular economy. Thus, connecting NAAT to a glucometer readout output provides a potential for low-cost, portable nucleic acid detection method that meets PoC needs along with environmental responsibility^2,3^.

Glucometer based biosensors have been extensively applied for detecting multitudes of analytical targets. Invertase- and amylase-based systems have been immensely studied: functional DNA probes trigger release of an enzyme which converts sucrose or starch to glucose which is readable by a glucose meter^4,5^. As originally demonstrated by Lan et al., a DNA–invertase conjugate can be released upon the target binding and hydrolyze sucrose into glucose, enabling a non-glucose target identification^6^. Similarly, DNA-functionalized hydrogels can trap glucoamylase (an amylose-to-glucose enzyme). Upon target-induced aptamer binding, the hydrogel then collapses and triggers glucoamylase release, converting amylose to glucose, thus enabling glucometer readout^7^. These approaches cleverly translate the binding events into output glucose signals. More recently, G-quadruplex DNAzymes have been used as signal transducers: in one scheme, a hemin/G4 DNAzyme catalyzes NADH oxidation, causing a drop in glucometer current^6^. Such DNAzymes has been integrated with CRISPR-Cas amplification for glucometer based detection of nucleic acid targets^6^. In another, hemin/G4 DNAzyme has been utilized to detect dengue virus nucleic acid sequence via a glucometer readout^8^.

However, all these “repurposed” glucometer methods have drawbacks. Invertase- or amylase-based sensors have the need for conjugating large enzymes to probes or the of requirement of co-expressing reporter enzymes, thereby appending complexity and higher costs. Hydrogels and nanocarriers must be precisely prepared for optimal functionality and robust performance^9^. DNAzyme methods avoid proteins but depend on particularly expensive reagents: for instance, many DNAzyme systems rely on costly cofactors like NADH or hemin^6^ (requiring fresh NAD^+^ reduction), which compromises their affordability and accessibility. Taken together, conventional and traditional enzyme-linked glucometer assays often require elaborate sample preparation and reagents, limiting their true “instrument-free” operation and hence are not feasible for PoC detection at remote areas.

We propose an alternative: exploiting the ferricyanide/ferrocyanide mediator chemistry which is at the core of glucometers^10^. In typical glucose oxidase based glucometer devices, glucose is oxidized to gluconic acid, which in turn reduces [Fe(CN)_6_]^3−^ to [Fe(CN)_6_]4−. The resultant ferrocyanide is then re-oxidized at the glucometer test strip electrode to produce the meter’s current^10^. Accordingly, the meter simply measures the redox state of the ferricyanide/ferrocyanide couple. Notably, any chemical agent that alters this redox pair will lead to the trigger of the same current. For example, ascorbic acid - generated by alkaline phosphatase acting on the ascorbic acid-phosphate, readily reduces K_3_[Fe(CN)_6_] to K_4_[Fe(CN)_6_], producing a glucometer signal^11^.

This suggests that a simple principle of non-electrocatalytic signal transduction through the ferricyanide/ferrocyanide redox couple could enable us to detect the presence of nucleic acid/NAAT product via the glucometer readout. In such schemes, a NAAT reaction, either directly or indirectly converting K_3_[Fe(CN)_6_] to K_4_[Fe(CN)_6_] in proportion to nucleic acid concentration would trigger a glucometer signal. The glucometer would then read out a current directly tied to nucleic acid levels. This ferricyanide/ferrocyanide based approach avoids the use of any extra enzymes or labeling. The glucometer itself becomes the detector for nucleic acid triggered redox changes. In summary, by exploiting the intrinsic [Fe(CN)_6_]^3−^/[Fe(CN) _6_]^4−^ chemistry of glucometers, we can repurpose them into simple, low-cost nucleic acid detection platforms.

Our research seeks to elucidate, that the presence and absence of DNA can be detected using a simple glucometer via the methylene blue (MB)/leucomethylene blue (LMB) mediated redox interconversion of potassium ferricyanide and ferrocyanide. We hypothesized that presence and subsequent intercalation of MB into DNA would influence the former’s interaction with ferrocyanide/ferricyanide redox couple, which in turn would influence glucometer readings. Therefore, the presence/absence of DNA (and by extension that of NAAT) and the dichotomy between the MB/LMB and ferricyanide/ferrocyanide dual redox couples can be explicitly detected via the glucometer signal readout. Through ratiometric absorbance analysis, we characterized the significance of buffer pH, salt conditions, external heat stimulus, concentrations of potassium ferricyanide, methylene blue, buffer and adjuvant to achieve optimal sensitivity for nucleic acid sensing. Subsequently, integration of aforementioned parameters into PCR, RCA, and LAMP were performed for investigating the assay’s compatibility with NAATs towards obtaining differential glucometer readouts enabling precise detection. Together these findings, would position a ubiquitous glucometer into a versatile nucleic acid and NAAT detection platform for scalable, sustainable and decentralized diagnostics.

## Materials and Methods

Please see Supplementary Material

## Results

### Design of experiments

The design of experiments followed a proleptic rationale based on the hypothesis that the presence and absence of DNA can be inferred by leveraging MB/LMB mediated redox dynamics of ferricyanide/ferrocyanide (Figure 1A). Furthermore, it was critical to evaluate whether this interaction would yield a significant readout change in the glucometer readings. It was also essential to identify the suitable buffer pH, optimal ferricyanide and MB concentration, and signal enhancement effects of adjuvant. Additionally, a crucial aspect of the design was also to ensure performance compatibility of the assay with NAATs such as PCR, RCA and LAMP. To systematically investigate these variables, we began by optimizing the pH of the buffer, effect of salt and heating conditions (Figure 1B). In subsequent experiments, we aimed to optimize the concentration of ferricyanide to identify an optimal operational concentration for differential detection (Figure 1B). Next, reactions with range of concentrations of MB were employed to evaluate the amount suitable for optimal sensitivity for detection of presence and absence of DNA (Figure 1B). To accentuate the signal difference between (+)DNA and (-)DNA conditions, an adjuvant was then utilized and optimized by studying a range of concentrations to achieve high sensitivity (Figure 1B). Finally, the assay was further studied for its compatibility and feasibility with NAATs integrated with differential glucometer readout (Figure 1B). In order to obtain the optimal assay conditions, our study adopted a synergistic combination of ferricyanide’s, and MB’s selective spectrophotometric responses (405 and 595 nm, respectively) coupled with glucometer measurement (Scheme 1A). Concurrently, we confirmed that potassium ferrocyanide was undetectable by absorbance at 405nm or by glucometer (giving a “Lo” signal, indicating a sample concentration below its limit of detection).

**Figure 1.**
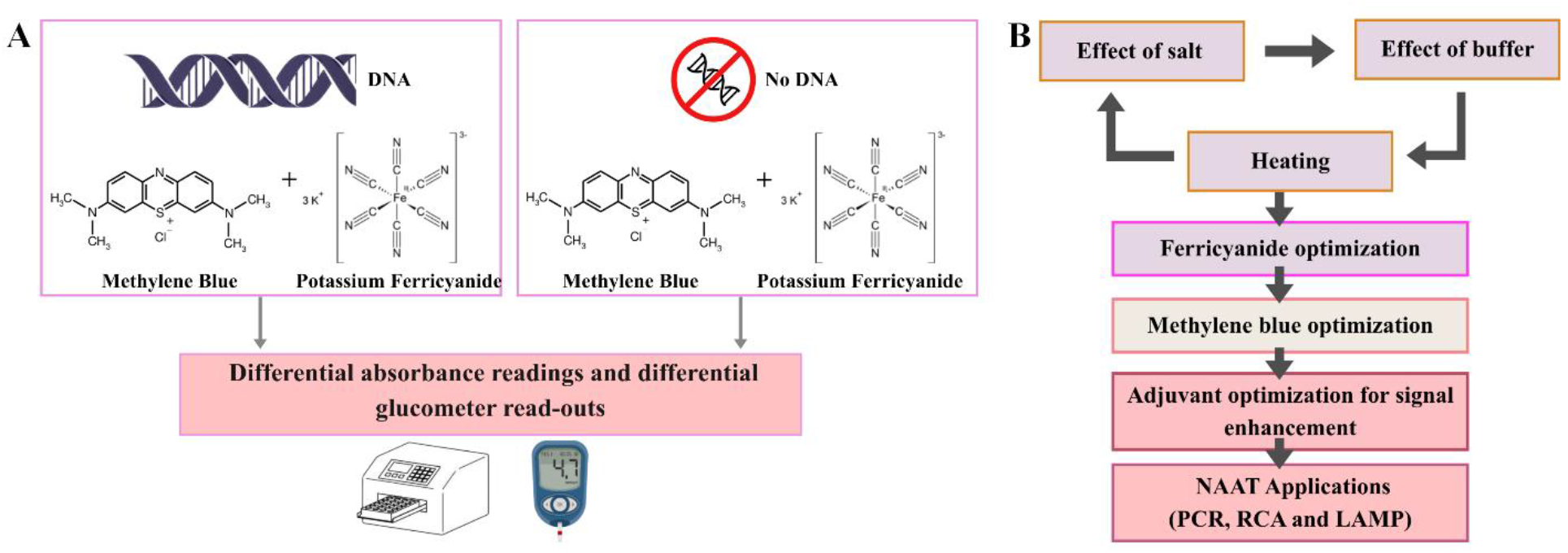
Schematic representation of the functional principle and experimental framework of the study. A, Illustration of the key reaction analytes and the underlying principle of detection of DNA using spectroscopic signatures and glucometer readouts. B, Experimental workflow depicting the effects of assay conditions (salt, buffer and heat), ferricyanide optimization, methylene blue optimization, optimization of adjuvant for signal enhancement, and application of the optimized techniques for various NAATs used in the study.

The working of glucometer was employed for quantitative differential detection of presence and absence of DNA. The standard glucometer strip has an enzyme layer, working electrodes and reference electrode (Figure 2A). The reaction analyte (1 µL-2 µL) droplet when brought in contact with the glucometer strip undergoes a biochemical reaction (Figure 2A). The enzyme layer contains glucose oxidase, which oxidizes the glucose to gluconic acid. The reduced enzyme then reacts with the potassium ferricyanide to reduce it to potassium ferrocyanide (Figure 2A). The resultant ferrocyanide loses its electrons to convert back to ferricyanide in the glucometer due to small, applied voltage at the working electrode. This specific feature of ferricyanide is utilized to detect DNA in tandem with the help of MB-LMB redox couple (Figure 2B).

**Figure 2.**
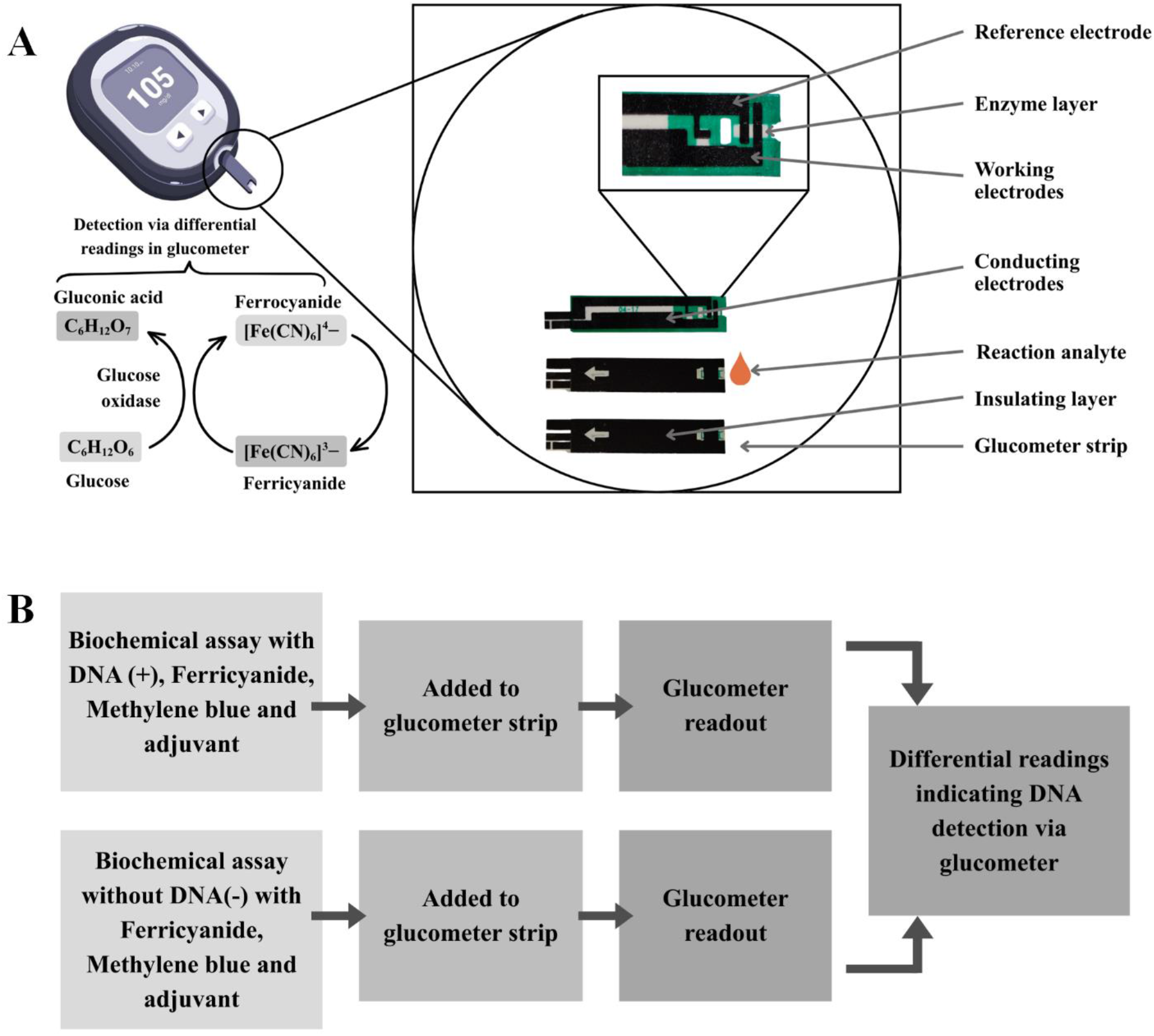
Glucometer architecture and assay readout workflow. A, Structural components of the glucometer device and glucometer strip depicting the underlying principle of electrochemical signal generation. It illustrates the glucose-mediated electrochemical signal transduction through enzymatic oxidation of glucose to gluconic acid coupled with ferricyanide/ferrocyanide redox system. B, Schematics representing the workflow of the assay measurement (DNA detection) using the glucometer, outlining the variable parameters (differential conditions), reaction analytes and signal readout for quantitative analysis.

### Hypothesis validation and optimization of buffer pH

We hypothesize that ferricyanide-mediated signal transduction via the effect of MB-LMB redox reaction can be used to detect presence and absence of nucleic acid (DNA) using a glucometer (Figure 2A). This stems from the fact that MB’s intercalation into DNA attenuates the former’s electron transfer ability during electrochemical sensing^12^. Analysis was done to identify the optimum pH of the buffer Tris-HCl under which the assay exhibited the maximal signal difference in DNA-present and DNA-absent conditions. Thereby, we investigated the buffer Tris-HCl at different pH values such as 7.0, 7.6, 8.0 and 8.5; wherein ferricyanide absorbance measurement was carried out at 405 nm (for ferricyanide) and 595 nm (for methylene blue) for a time course study (0-30 min) (Figure 3A). Further, we wanted to analyze the effect of presence of salt and external heating stimulus. These experiments were performed in presence and absence of DNA (referred to as “(+) DNA” and “with DNA”, or “(-) DNA” and “without DNA”, respectively) and represented in terms of ratiometric absorbance (RA) (A_405_/A_595_) at 5 minutes and 30 minutes (Figure 3A). While the time course study helped us understand the dependence of time on the signal transduction, the difference in the RA indicated whether the assay could distinguish between DNA presence and absence. MB binding to DNA causes former’s absorbance to decrease, resulting into decrease in RA^13^. The difference in RA between (+)DNA and (-)DNA conditions would thus correspond to improved and quantitative differentiation of the presence of DNA from its absence.

**Figure 3.**
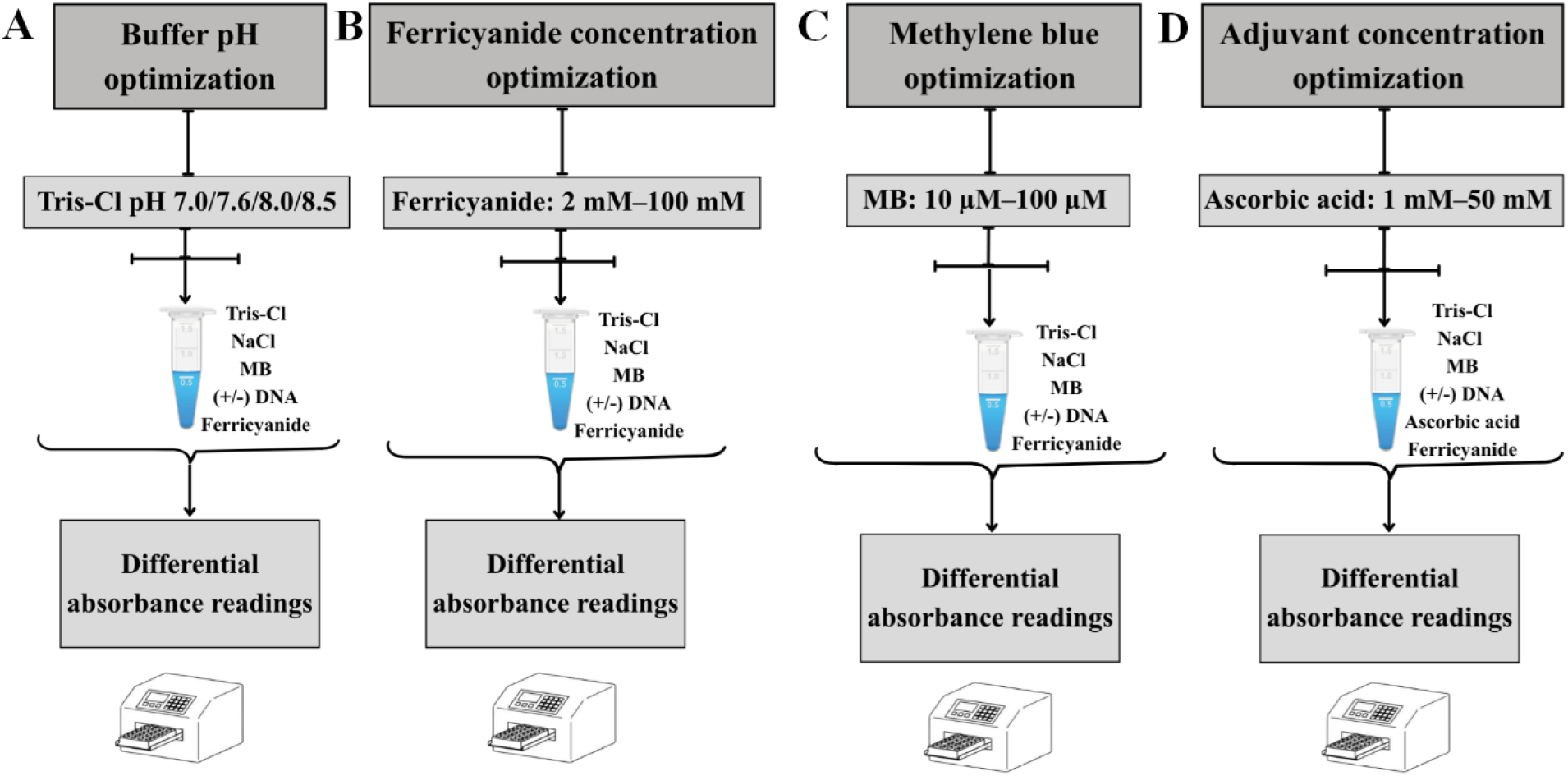
Various optimization conditions utilized in this study. Panel A, Schematics of proof-of-concept and buffer pH optimization for tris-HCl buffers of pH 7.0, 7.6, 8.0, 8.5 (Panel A), potassium ferricyanide (2 – 100 mM) optimization (Panel B), methylene blue (10 – 100 μM) optimization (Panel C), and adjuvant (ascorbic acid, 1 – 50 mM) optimization (Panel D). Experiments were performed in presence or absence of DNA (25 ng/μL) with measurement of ratiometric absorbance (A_405_/A_595_) values.

All conditions and pH variations exhibited an absorbance difference in the presence and absence of DNA (Figure 4A-D). A consistent trend that emerged across all tested conditions here as well as in later studies, was that the RA of “DNA-present” was significantly higher compared with the RA of “DNA-absent” condition. The trend exhibited a slight decrease of RA over the time, indicating time dependent signal response property of the assay. The assay yielded a significant difference in presence and absence of DNA conditions at pH 7.0 and pH 7.6 and thereby showed pH dependence property of the assay (Figure 4A and B).

**Figure 4.**
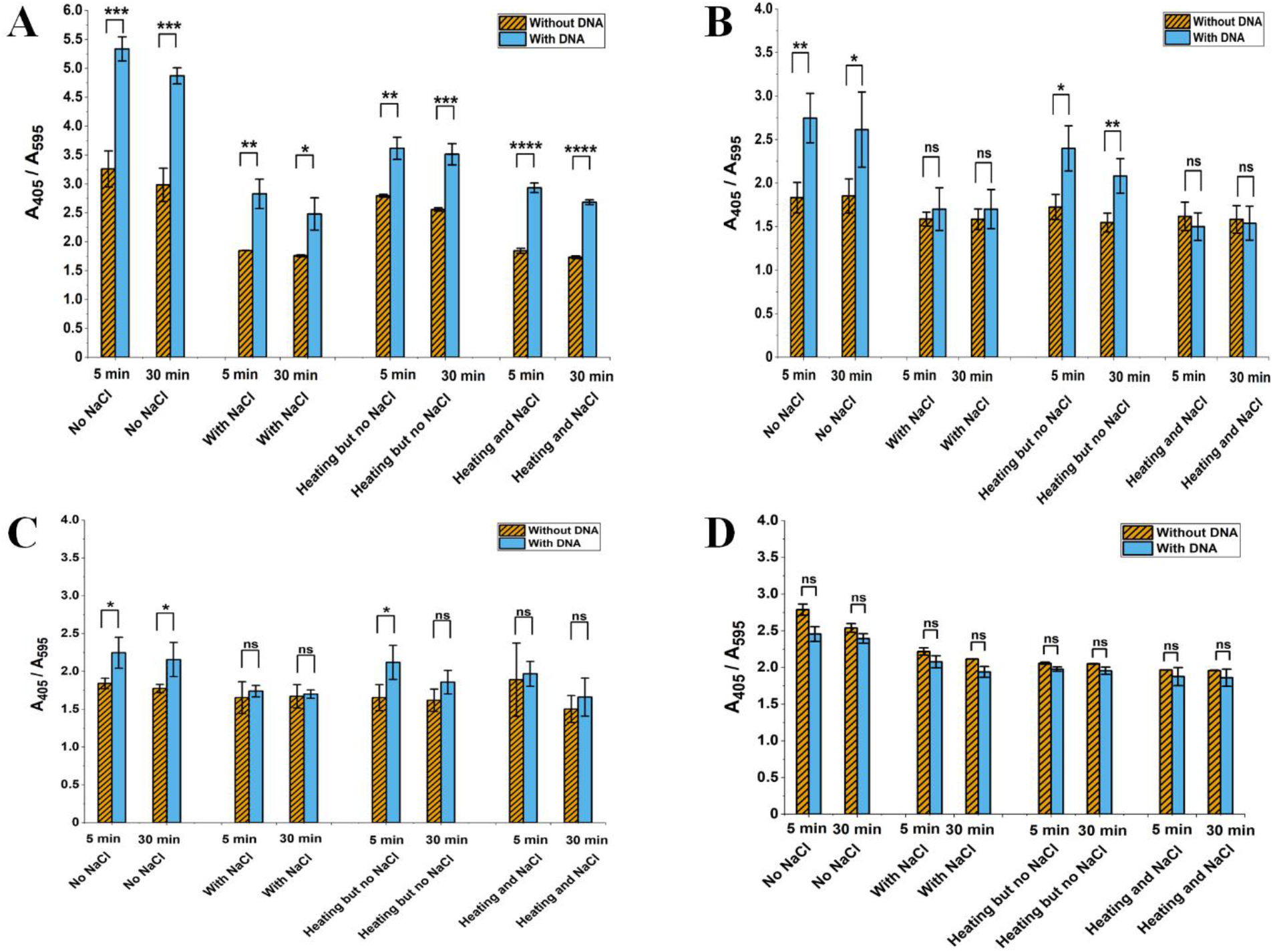
Optimization of buffer pH and assay conditions for optimum difference between (+) DNA (25 ng/μL) and (−) DNA conditions based on Figure 3A schematics. Panel A-D represents the ratiometric absorbance (A_405_/A_595_) with combination effects of varying salt and heating conditions at pH 7.0 (Panel A), pH 7.6 (panel B), pH 8.0 (Panel C), and pH 8.5 (panel D) in presence and absence of DNA at 5 and 30 minutes time points. Experiments were carried out with 2 mM potassium ferricyanide and 80 μM MB. Error bars represent standard deviation (n = 4). Statistical analysis performed using Student’s t-test. P < 0.05 (*), P < 0.01 (**), P < 0.001 (***), P < 0.0001 (****).

NAAT reactions such as PCR and LAMP utilizes heating and monovalent cations to partially denature DNA and stabilize duplex formation, respectively. Consequently, our assay probed the impact of presence of salt and external heating stimulus on the overall RA profile in the reaction assay. Heating causes DNA to denature, while the presence of salt (NaCl) the DNA tightens the duplex structure. Therefore, presence of NaCl enables lesser intercalation of MB into the DNA backbone, resulting lesser hypochromic change in MB absorbance, and lesser RA magnitude. On the other hand, heating temporarily untangles the DNA, causing MB to better intercalate into DNA, resulting in a greater hypochromic change in MB absorbance, and subsequently greater RA magnitude. These trends were clearly reflected in the RA profiles of various Tris-HCl buffer variation studies, predominantly so in pH 7.0 and 7.6. On the other hand, the RA profiles for Tris-HCl pH 8.0 and 8.5 could not yield significant difference between DNA presence and absence, underscoring the effect of pH on the assay.

Overall, these spectroscopic experiments supported our novel hypothesis that DNA can be detected using the interaction between DNA, MB, and ferricyanide. The greater difference in DNA-present and DNA-absent conditions at pH 7.0 and pH 7.6 suggested that they provide a more suitable buffering environment for the assay compared to pH 8.0 and pH 8.5. Furthermore, presence of monovalent cation and heating, simulating a NAAT-like environment, was capable of differentiating the DNA presence/absence as well, thus further corroborating our experimental optimization. Due to reduced difference in DNA-present/DNA-absent conditions, we pursued with Tris-HCl pH 7.0 and pH 7.6 for our further experiments.

### Optimization of potassium ferricyanide concentration

Subsequent to the evaluation of the buffer pH suitability, salt, and heating conditions, we next determined the optimal concentration of potassium ferricyanide for better differentiating the presence and absence of DNA (Figure 3B). We observed a greater difference in RA at 40 mM and 100 mM ferricyanide concentrations between the DNA-present and DNA-absent conditions in both Tris-HCl pH 7.0 and 7.6 (Figure 5A and B). While at lower concentrations (2 mM, 6 mM and 18 mM) the difference in RA for the presence and absence of DNA remained consistently lower for pH 7.0 and pH 7.6 at both time points. In DNA-present condition, the MB would be intercalated into DNA, causing hypochromic MB absorbance, resulting in a higher RA compared to the DNA-absent condition. Overall, these results established that 40 mM concentration of ferricyanide is the most optimum for a significant distinction between presence and absence of DNA.

**Figure 5.**
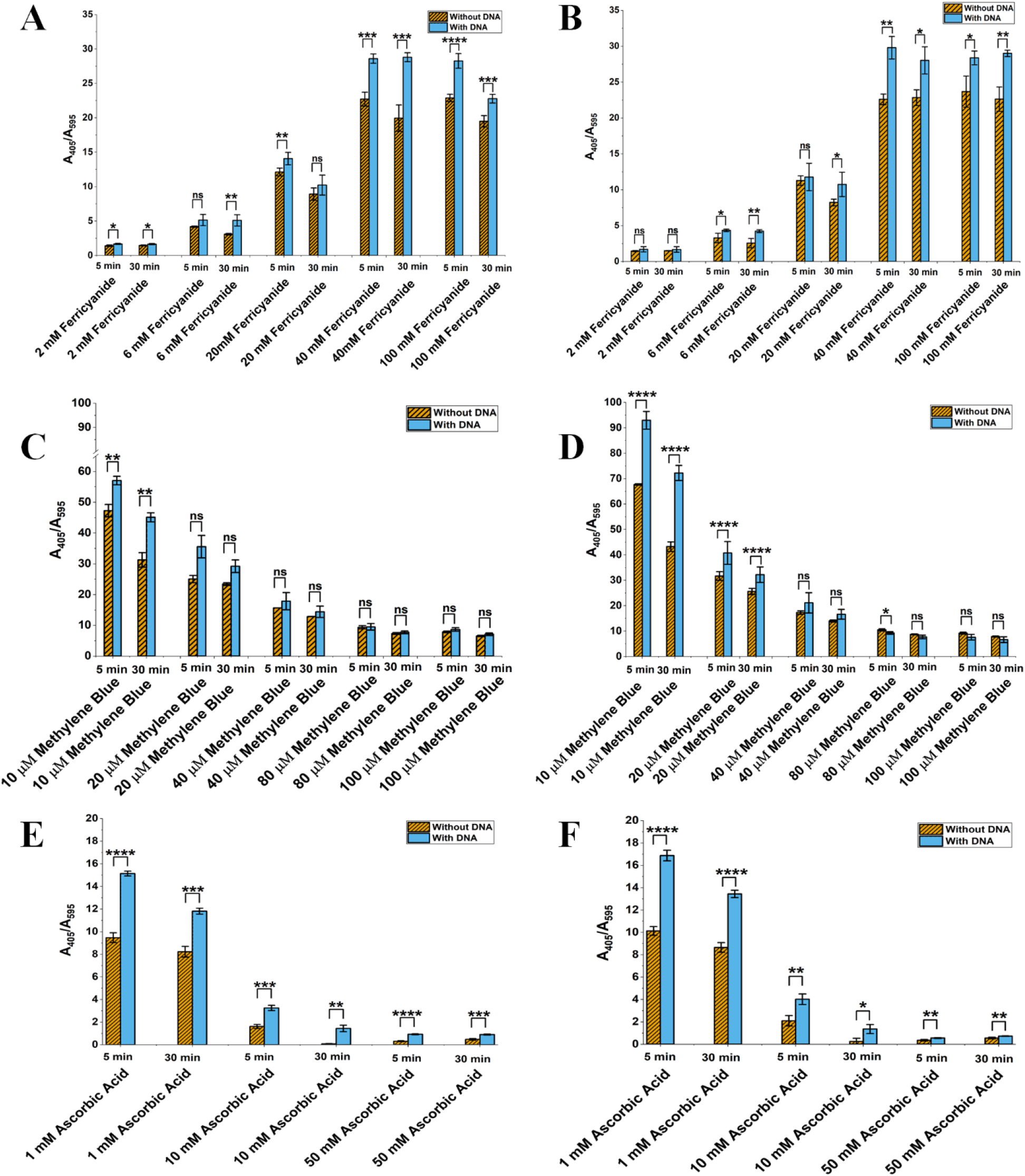
Ratiometric absorbance-based optimization of potassium ferricyanide, MB, and ascorbic acid (adjuvant) concentrations for differentiating (+) DNA (25 ng/μL) and (−) DNA. Experiments were performed based on Figure 3B-D schematics. A and B, Ratiometric absorbance (A_405_/A_595_) with varying ferricyanide concentrations (2 mM – 100 mM), 80 μM MB in presence/absence of DNA at 5 and 30 minutes at tris-HCl pH 7.0 (Panel A) and tris-HCl pH 7.6 (Panel B). C and D, Ratiometric absorbance (A_405_/A_595_) with varying MB concentrations (10– 100 μM), 20 mM potassium ferricyanide in presence/absence of DNA at 5 and 30 minutes at tris-HCl pH 7.0 (Panel C) and tris-HCl pH 7.6 (Panel D). E and F, Ratiometric absorbance (A_405_/A_595_) with varying ascorbic acid (adjuvant) concentrations (1 – 50 mM), 20 mM potassium ferricyanide, 10 μM MB in presence/absence of DNA at 5 and 30 minutes at tris-HCl pH 7.0 (Panel E) and tris-HCl pH 7.6 (Panel F). Error bars represent standard deviation (n = 4). Statistical analysis performed using Student’s t-test. P < 0.05 (*), P < 0.01 (**), P < 0.001 (***), P < 0.0001 (****).

### Optimization of MB concentration

Following the assessment of optimum concentration of ferricyanide, we next evaluated the optimal concentration of MB to further improve signal difference between presence and absence of DNA (Figure 3C). We carried out the assessment using a range of methylene blue concentrations: 10, 20, 40, 80 and 100 µM. We found that 10 µM MB would be the most appropriate concentration to go ahead due to having higher difference in RA between the DNA-present and DNA-absent conditions in both Tris-HCl pH 7.0 and 7.6 (Figure 5C and D). Collectively, these results substantiate that 10 µM concentration of MB is the most optimum for a significant distinction between presence and absence of DNA in subsequent studies.

### Optimization of ascorbic acid concentration

Subsequent to the evaluation of optimal buffer pH, salt and heating conditions, optimal concentrations of ferricyanide and methylene blue, we next determined the optimum concentration of ascorbic acid as an adjuvant in this reaction (Figure 3D). Since ascorbic acid reduces Fe(III) to Fe(II)^14^, we hypothesized that presence of ascorbic acid as an adjuvant would lead to greater signal difference between DNA presence/absence. Accordingly, the assessment was carried out using a range of ascorbic acid concentrations: 1, 10 and 50 mM. Higher ascorbic acid concentrations resulted into lower RA values (A_405_/A_595_) owing to reduction of ferricyanide (Fe(III)) (A_405_ nm), thus validating our hypothesis (Figure 5E and F). We observed a greater difference in RA at 1 mM and 10 mM ascorbic acid concentrations between the DNA-present and DNA-absent conditions in both Tris-HCl pH 7.0 and 7.6. Notably, 10 mM ascorbic acid yielded the highest relative difference in RA (∼2 – 20 fold) between the DNA-present and DNA-absent conditions in both Tris-HCl pH 7.0 and 7.6. In comparison 1 and 50 mM ascorbic acid only provided ∼1.2 – 1.8 fold RA difference between presence/absence of DNA. Hence, 10 mM ascorbic acid, owing to its significant distinction between presence and absence of DNA, was utilized in subsequent analyses.

### Compatibility with NAATs and enhanced detection of PCR product via glucometer

After the evaluation of optimal buffer pH, salt, heating conditions, ferricyanide, MB and adjuvant concentrations, we next scrutinized the compatibility of our assay with NAATs: PCR, RCA and LAMP integrated with glucometer signal readouts. The PCR was carried out in presence of MB and then supplemented with ferricyanide as well as adjuvant, followed by glucometer measurements over a time course of 30 minutes (Figure 6A). While probing the assay with PCR, we also analysed the signal enhancing effect of adjuvant on the glucometer signal readout. We found that PCR led to an amplified signal difference between DNA presence/absence conditions which was further enhanced when adjuvant was supplemented (Figure 6B).

**Figure 6.**
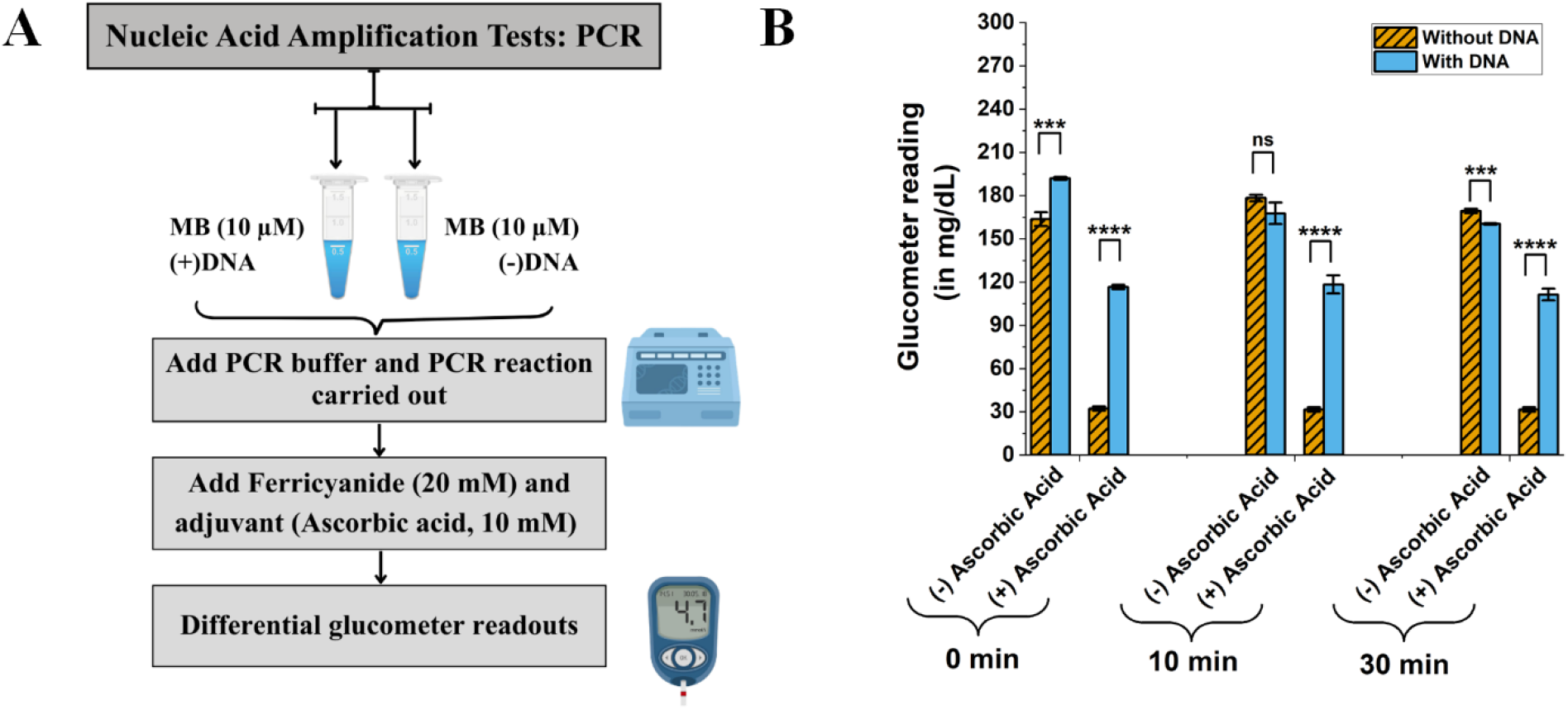
Effect of ascorbic acid on glucometer-based assessment of PCR reaction (with and without DNA template) for detecting *E. coli* gDNA using optimized MB (10 μM MB), potassium ferricyanide (20 mM), and ascorbic acid (10 mM). A, Schematics of experiments. B, Time point (0, 10, 30 min) analysis of glucometer response for PCR reaction on 10^6^ copies/reaction of *E. coli* gDNA template as a function of ascorbic acid presence. Measurements were taken after 0 min, 10 min, or 30 min after ferricyanide addition. Error bars represent standard deviation (n = 4). Statistical analysis performed using Student’s t-test. P < 0.05 (*), P < 0.01 (**), P < 0.001 (***), P < 0.0001 (****).

### Glucometer-enabled limit of detection (LOD) of PCR

Subsequent to the evaluation of optimal buffer pH, salt, heating conditions, ferricyanide, MB, adjuvant concentrations and compatibility with PCR assay and temperatures, we next determined the glucometer-based LOD for our developed assay and pipeline. The assessment was performed using a negative control (DNA(-)) and a range of DNA copies – 10^0^, 10^1^, 10^2^, 10^3^, 10^4^, 10^5^, 10^6^, 10^7^ and 10^8^. In addition, the assay also measured the glucometer signal magnitudes over a time period of 30 min. The recorded glucometer readings showed differential readings signifying the presence and absence of DNA conditions over a varied range of DNA copy numbers for 0, 10, and 30 min time points (Figure 7A, B, and C). The differential glucometer readings and R-square values (>0.9) demonstrated that the assay has a broad linear dynamic range and not just circumscribed to single type of copy number (Figure 7D and E). The LOD for the glucometer-enabled PCR measurement was 10^2^ copies/reaction, which reflected remarkable assay performance with sensitivity at par with reported fluorescence- and electrochemistry-based NAAT readouts^15,16^. On the contrary, when LOD assay was performed with 80 µM MB (without altering previously optimized assay conditions), we failed to obtain LOD below 10^4^ copies/reaction. The better LOD performance of 10 µM over 80 µM further confirmed the success of our assay optimization workflow (Supplementary Material Figure S1). Moreover, relative standard deviation (RSD) magnitude measured for both 10^4^ and 10^8^ copies across all three time points were below ∼10%, thereby proving the assay’s high degree of reproducibility (Figure 8). In summary, these experiments demonstrated that our optimal assay conditions can be harnessed for glucometer-enabled reproducible detection of PCR reaction products over a broad dynamic range, thereby validating the assay’s resilient performance and exceptional sensitivity.

**Figure 7.**
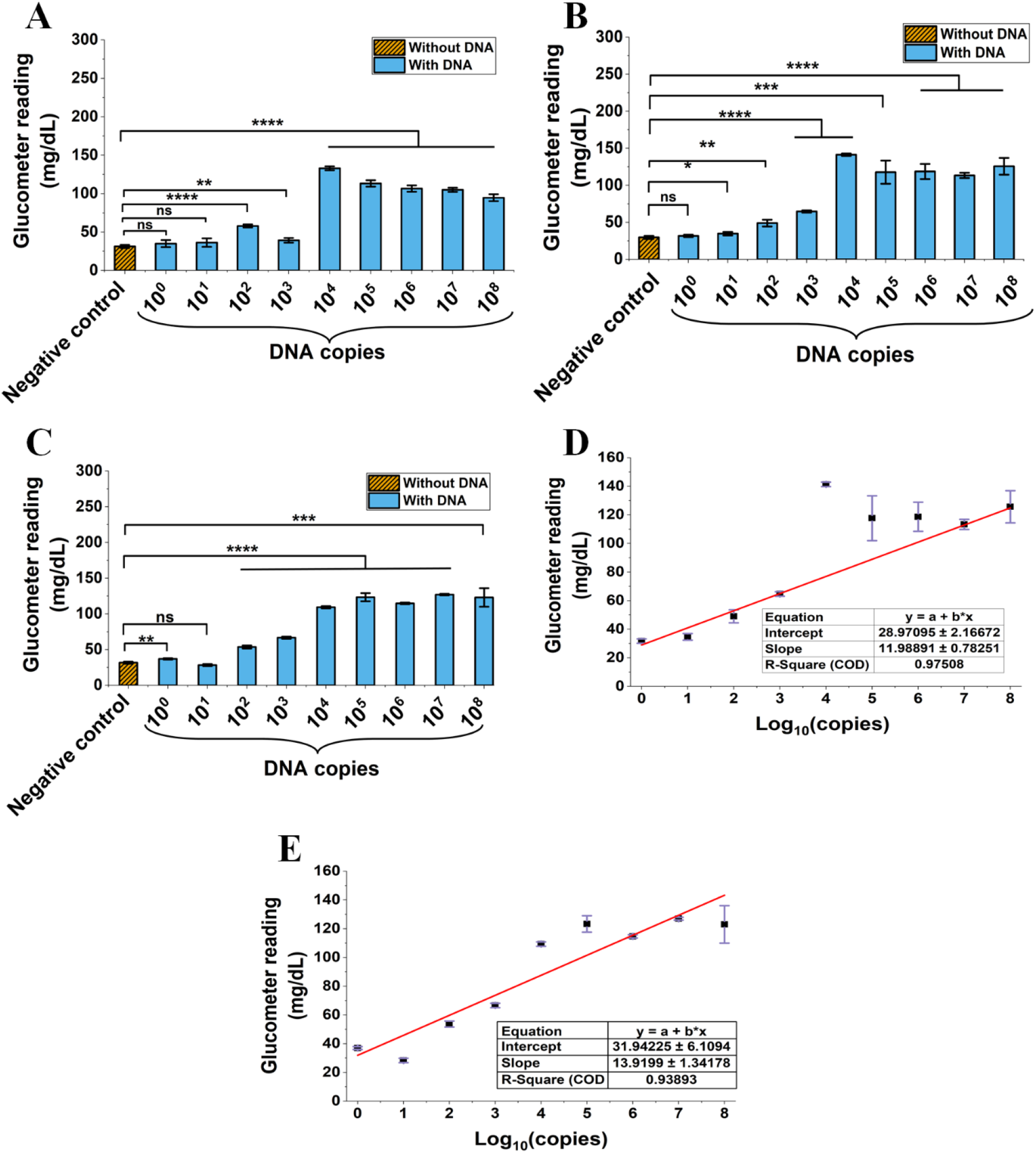
LOD and reproducibility assessment for PCR-based detection of *E. coli* gDNA. A – C, Glucometer response for detecting 10^0^ – 10^8^ copies/reaction of *E. coli* gDNA for measurements taken after 0 min (panel A), 10 min (panel B), and 30 min (panel C) of ferricyanide addition. D and E, Linear fit for LOD data for 10 min (Panel D) and 30 min (Panel E) PCR measurements. The glucometer reading at 10^4^ copies for 10 min measurement (Panel D) were not taken into account for the linear fitting. Optimized concentrations of MB (10 μM), potassium ferricyanide (20 mM), and ascorbic acid (10 mM) were used for these experiments. Error bars represent standard deviation (n = 4). Statistical analysis performed using Student’s t-test. P < 0.05 (*), P < 0.01 (**), P < 0.001 (***), P < 0.0001 (****).

**Figure 8.**
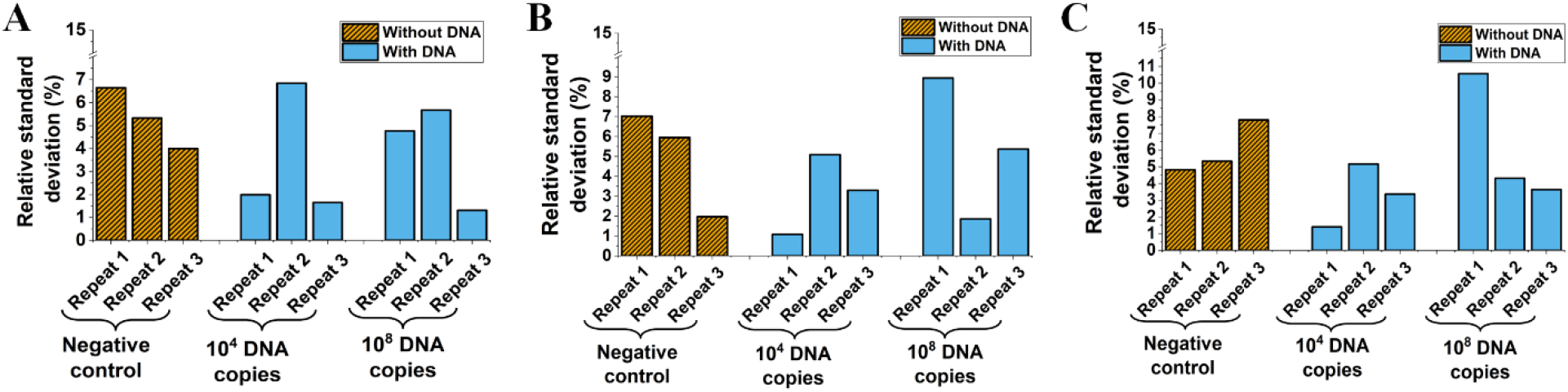
Reproducibility of glucometer-enabled PCR assay. A – C, RSD values for negative control, 10^4^ copies/reaction, 10^8^ copies/reaction for three independent PCR repeats for glucometer measurements taken after 0 min (panel A), 10 min (panel B), and 30 min (panel C) of ferricyanide addition.

### Detection of RCA amplicon via glucometer readout

Next, we checked the assay compatibility with the RCA using dengue virus serotype 2 R3 polyprotein sequence as the target. The circular DNA was synthesized using splint padlock ligation followed by RCA initiated through use of target as the primer. The amplification step was caried out in presence of MB while ferricyanide and ascorbic acid were being added post the RCA experimental procedure. The glucometer readouts were recorded for a course of 30 minutes (Figure 9A). As shown in Figure 9B, a significantly large difference between DNA-present and DNA-absent conditions was observed, where the DNA-present condition gave a low glucometer reading. Despite having a different salt and buffer conditions than that of PCR, our results clearly showed that the novel assay pipeline developed is compatible with RCA as well.

**Figure 9.**
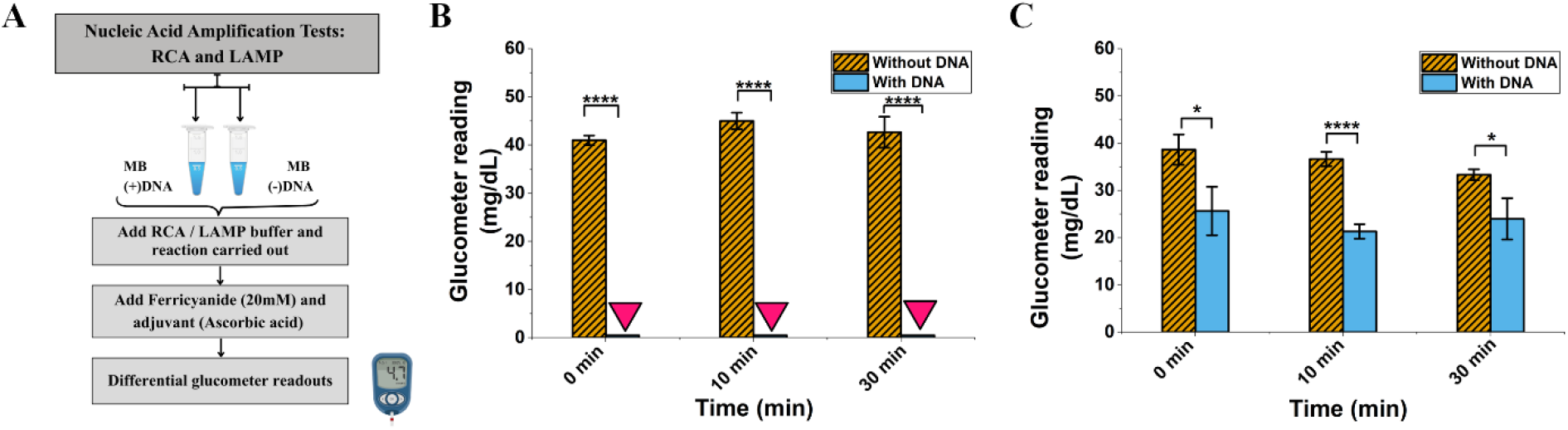
Glucometer-enabled detection of RCA and LAMP assays using optimized conditions. A, Schematics of assay. B, Glucometer response for HRCA-mediated detection of dengue virus serotype-2 biomarker sequence (1 nM) measured at 0, 10, 30 min after ferricyanide addition. B, Glucometer response for LAMP-mediated detection of SARS-CoV-2 RdRp sequence (10^6^ copies/reaction) measured at 0, 10, 30 min after ferricyanide addition. Optimized concentrations of MB (10 μM), potassium ferricyanide (20 mM), and ascorbic acid (10 mM) were used for these experiments. Error bars represent standard deviation (n = 4). Statistical analysis performed using Student’s t-test. P < 0.05 (*), P < 0.01 (**), P < 0.001 (***), P < 0.0001 (****).

### Detection of LAMP amplicon via glucometer readout

Next the LAMP assay was performed with SARS-CoV-2 RdRp gene present in a plasmid as the molecular target. LAMP was executed in presence of MB while ferricyanide and ascorbic acid were added post the LAMP experimental procedure. The glucometer readings were being documented for a course of 30 minutes (Figure 9A). As shown in Figure 9B, a significant difference (∼2 fold) between DNA-presence and absence conditions was noted. Similar to RCA and unlike PCR, the glucometer readouts in DNA-present conditions were of lesser magnitude compared to DNA-absent conditions. It was plausibly due to generation of concatenated amplicons in both RCA and LAMP that might have encapsulated MB and ferricyanide, hindering the redox chemistry^17,18^. In summary, the results showed that the novel assay pipeline developed is compatible with the LAMP assay, where the signal amplification effects of LAMP were detectable through the enhanced differential glucometer readouts.

## Discussion

Our novel glucometer-based nucleic acid and NAAT sensor meets the key needs identified in our research: it is low-cost, label-free, portable, and highly sensitive. By replacing the conventional expensive lab potentiostats, electrodes, and qRT-PCR, our assay leverages an off-the-shelf inexpensive glucose meter (INR 500 – 1500, ∼ USD 5 – 15 per unit) without sacrificing any detection limits. Indeed, sensitivity down to hundreds of copies matches and sometimes exceeds previously reported glucometer nucleic acid assays^4,19,20^, validating that the glucometer signal can robustly quantify the amplified nucleic acids. Unlike typical electrochemical DNA sensors, no specific enzymes (such as invertase^4^) or any sophisticated labels (e.g., ferrocene^21^) are added in the assay to obtain the final readout. The inherent ferricyanide/ferrocyanide reaction in the meter suffices to generate a significant signal difference between the presence and absence of nucleic acid in the form of amplicon. This simplicity is a major and key novelty: we established a general platform where any NAAT output (PCR, RCA, LAMP) is directly translated into glucometer readout signals.

Our research work thus fills the gap of “equipment-free” nucleic acid detection. Beyond cost, convenience and simplicity, the approach also offers various sustainability advantages. Most personal glucometers already exist in vast numbers at any pharmacy worldwide. Repurposing the glucometers prevents new device manufacturing, aligning with a sustainable circular-economy approach. In contrast to traditional nucleic acid detection methods that require centralized and energy-intensive instrumentation such as qRT-PCR and potentiostats, the glucometer-based platform supports a low-infrastructure analytical workflow with reduced operational and energy demands. Additionally, the label-free assay uses small reaction volumes and cost-effective redox mediators, eliminating the need for fluorescent labels, enzymatic conjugates, and complex probe modifications. This approach reduces reagent complexity, chemical waste, and overall resource consumption. The glucometer is one of the widely available, inexpensive POCT device with excellent ergonomic design; enabling reproducible and quantitative readouts^10,20,22^. At limited-resource settings and remote locations, the glucometer’s design – intuitive interface and high battery power-makes it durable, portable and user-friendly for non-experts. In practice, a health worker in the field could add a few drops of amplified sample to a test strip and quickly read the result on the familiar glucometer, without any specialized training or expertise. Our platform capitalizes on these attributes and key USPs.

In summary, our glucometer-based nucleic acid and NAAT detection platform provides a compact, label-free alternative to bench-top assays. It answers the urgent need for a democratized diagnostics: affordable, point-of-care testing and screening accessible outside the advanced lab at remote places. Our results highlight that coupling NAAT with a ubiquitous glucometer consumer device can dramatically widen the test deployment. Future work will consist of further optimization assays for reagent stability and integration (for example a self-contained cartridge with lyophilized reagents), and/or validate the device and assay pipeline on real clinical samples. Conclusively, this research and strategy paves the way for home or PoC field diagnostics of pathogens and biomarkers, thereby effectively bringing molecular tests into everyday healthcare.

## Conclusion

We demonstrated that a standard off-the-shelf glucometer can be repurposed into an ultra-low-cost, completely portable nucleic acid and NAAT detection assay. Our assay achieved a high sensitivity (comparable to generic qRT-PCR-based tests) while using only a simple and inexpensive glucometer with routine lab reagents. We systemically optimized experimental conditions such as buffer pH, salt, temperature, redox mediators, and adjuvant to achieve maximum signal difference in DNA(+/–). This resulted into an excellent LOD with the assay elucidating a linear to near-linear glucometer signal across a wide target range. We validated various NAAT workflows starting with standard PCR which yielded enhanced detectable signals for DNA within ∼ 1 h with 10^2^ copies/reaction as the LOD. Furthermore, isothermal NAATs such as RCA (operating at room temperature, assay time ∼2–3 h) and LAMP (operating at 60–65°C, assay time ∼1 h) also achieved significant differential sensitivity between DNA presence/absence conditions. Each method, despite their difference in salt, pH, and buffer parameters resulted into robust glucometer readings, confirming that varied amplification strategies are not just compatible with our novel assay but also capable of attaining differential signal readout. Fundamentally, unlike any conventional ELISA, fluorescence- or electrode-based methods, our assay platform is completely label-free and modified electrode free; the glucometer is both the transducer and readout in itself.

In essence, our glucometer-based NAAT integrates the ease of glucose testing with the utmost sensitivity of a qRT-PCR, at a fraction of the cost when compared to any existing technology. Given the glucometer’s global ubiquity and ergonomic design, our innovation promises to “democratize” molecular diagnostics, and hereby enabling rapid, on-site testing for infectious diseases and other genetic targets in any setting. By turning an everyday device into a general biosensor, this work has opened unprecedented opportunities for affordable point-of-care diagnostics and public health monitoring. By the integration of the advanced molecular diagnostic technique like PCR through an everyday inexpensive PoC device like glucometer, this strategy holds potential promise for a wide deployment in clinics, homes, resource-limited regions, and remote areas where access to healthcare is limited. Our results are on par and sometimes exceeds that from with the best reported glucometer-based nucleic acid detection assays while matching qRT-PCR sensitivity. Ultimately, transforming a glucometer into a nucleic acid and NAAT sensor would not only revolutionize the healthcare diagnostics but also democratize the point-of-care testing and public health monitoring and yet being sustainable and environmentally responsible.

## Abbreviations

NAAT: nucleic acid amplification test
PoC: point of care
POCT: point-of-care testing
MB: methylene blue
LMB: leucomethylene blue
PCR: polymerase chain reaction
RCA: rolling circle amplification
LAMP: loop-mediated isothermal amplification
RA: ratiometric absorbance

## Author Credit

AC, SP, SK, AT, MS, SG conceived the experiments and analyses. AC, SP, and SK performed the experiments. MS helped understood the biological aspects of nucleic acid amplification technology. AT helped understand the clinical aspect of the work. AC, SP, and SG performed the data analyses. AC, SP, and SG wrote the manuscripts. All authors read and approved the manuscript.

## Supplementary Information

Materials and methods with details experimental protocols; Additional Figures;

## Acknowledgement

This work was supported by the Indo-French Centre for the Promotion of Advanced Research (CEFIPRA, No. 70T09-2), IIIT-iHub grant (No. H4-006), DST INSPIRE PhD Fellowship to Ms Saba Parveen (No DST/INSPIRE/03/2021/002361), and Mahindra University internal funding to “Interdisciplinary Centre for Nanosensors and Nanomedicine”. The authors are grateful to Prof. Rajinder S. Chauhan, Prof. Bishnu Pal, and Prof Yajulu Medury (Vice-Chancellor of Mahindra University) for their encouragement and constant scholarly support.

