## Supplemental Data for "Transforming off-the-shelf personal glucose meter into a sustainable and decentralized label-free nucleic acid and NAAT detection platform"

### Materials and Methods

Potassium ferricyanide ($K_3[Fe(CN)_6]$) (cat# 59558), sodium chloride (NaCl) (cat# 33205), ascorbic acid ($C_6H_8O_6$) (cat# 32488) and methylene blue (cat# 31535) were purchased from SRL. Tris (hydroxymethyl) aminomethane (Tris) (cat# 252859) and ethylenediaminetetraacetic acid (EDTA) (cat# E5134) were purchased from Sigma-Aldrich. PCR master mix (cat# MBT061-100R) was purchased from HiMedia. E. coli genomic DNA was being used in the experiments. The spectrophotometer used was Thermo Scientific Multiskan FC. Personal glucometer such as Dr. Morepen GLuco One (model no: BG-03, license no: MFG/IVD2021/000034 and lot no: AIK237) and Apollo pharmacy blood glucose monitoring (model no: APG01, license no: MFG/IVD/2021/000034 and lot no: DIL272) and their manufacturer designated strips were purchased locally and was used as it for all analysis and experiments.

T4 polynucleotide kinase (PNK), T4 DNA ligase, ϕ29 DNA polymerase, dNTP,BSA and ϕ29 buffer were purchased from New England Biolabs, USA. Tris buffers, NaCl, spermidine, and DTT were purchased from Sisco Research Laboratories Pvt. Ltd. (SRL) -India. All solutions were prepared using ultrapure water obtained from a Millipore Milli-Q Type II water purification system. Phosphorylation, annealing, and ligation reactions were performed in an Eppendorf Mastercycler Nexus system. Real-time RCA studies were carried out in Bio-Rad CFX96 qRT-PCR and analysed using CFX Maestro software. Gel results were visualized and analysed by the Bio-Rad Gel documentation system. If necessary, oligonucleotide solutions were concentrated using Eppendorf Concentrator Plus speedvac at ambient temperature.

*Escherichia coli* strK-12 substr. MG1655 plasmid construct with RNA-dependent RNA polymerase (*RdRp*) gene with T7 RNA polymerase promoter (4538 bp) was purchased from Addgene (plasmid #14567, https://www.addgene.org/145671/)**.** The Bst 2.0 polymerase, RTx enzyme, dNTP, and SnaBI were procured from NEB, USA. Molecular biology grade water was purchased from HiMedia, India. The RNase inhibitor was purchased from Takara. Streptavidin-coated magnetic beads were purchased from Sigma-Aldrich (# 11641778001) or Invitrogen (Dynabeads M − 280). 5′-biotinylated probe having the b sequence (5’-[BIO]-AAA AAA AAA ACG AGC AAG AAC AAG TGA GGC CAT AAT TC, HPLC purified, $T_m$ = 57.6 °C at 50 mM NaCl and 0.25 μM oligonucleotide concentration) was purchased from Sigma-Aldrich. Primer oligonucleotides (desalting purified) were purchased from Eurofin or Sigma-Aldrich, India. LAMP primer set 2 sequences were as follows: F3 (CGA TAA GTA TGT CCG CAA TT), B3

(GCT TCA GAC ATA AAA ACA TTG T), FIP (ATG CGT AAA ACT CAT TCA CAA AGT CCA ACA CAGACT TTA TGA GTG TC), BIP (TGA TAC TCT CTG ACG ATG CTG TTT AAA GTT CTTTAT GCT AGC CAC) Loop F (TGT GTC AAC ATC TCT ATT TCT ATA G), Loop B (TCA ATA GCA CTT ATG CAT CTC AAG G)

All the experiments utilize type I water. All concentrations mentioned below are final concentrations in the actual reaction or process unless stated otherwise.

**Buffer pH optimization:** Tris-HCl (Tris(hydroxymethyl)aminomethane Hydrochloride) (final 20mM) was being used as the buffer for the reaction assay. The pH of the buffer was optimized by studying a range of pH values –7.0, 7.6, 8.0 and 8.5. For each pH four experimental conditions were being evaluated, one without salt, second with salt supplementation, third without salt and without heating, and lastly with salt and heating. All four experiments were done with two separate critical conditions: DNA present ((+)DNA) and DNA absent ((-)DNA). For each condition 70 µL (final) of reaction assay was being prepared in replicates, containing the Tris-HCl buffer (final 20 mM), +/-NaCl (final 50 mM), methylene blue (final 40 µM), and +/-DNA (15 ng/µL). The experiments with heating condition were being heated at 95°C for 5 mins using a heat block. 63 µL of all the assays were being then added to the 96-well plate in three or more replicates, and promptly 7 µL of potassium ferricyanide [$K_3Fe(CN)_6$] (final 2 mM) was added and gently mixed. Absorbances at 405 nm and 595 nm were being measured using the microplate reader. The kinetic loop program was set in the plate reader to measure absorbance readings at every 5-minute intervals, starting from 0 minutes and up until 30 minutes elapsed (i.e. at 0, 5, 10, 15, 20, 25 and 30 minutes), without the removal of the plate from the reader during the entire duration of absorbance measurement. The entirety of reaction was being performed at ambient room temperature apart from the heating condition. The recorded absorbance readings showed difference in presence and absence of DNA conditions. This difference reflected the presence/ absence of DNA and pH effects on the detection.

**Optimization of potassium ferricyanide concentration:** To determine the optimal potassium ferricyanide concentrations for maximizing the glucometer signal difference in the presence and absence of DNA. Varied concentrations of ferricyanide (2 mM, 6 mM, 18 mM, 40 mM and 100 mM) were evaluated in Tris-HCl buffer at pH 7.0 and pH 7.6, supplemented with 50 mM NaCl. All assays were conducted under the previously optimized experimental conditions (with heating).

For each ferricyanide concentration, reaction assays were prepared containing Tris-HCl buffer with a final concentration of 20 mM, NaCl 50 mM, methylene blue 40 µM, and DNA presence or absence with 25 ng/µL. Prior to ferricyanide addition, the reaction assays were heated at 95°C for 5 min followed by the immediate addition of potassium ferricyanide [$K_3Fe(CN)_6$] to achieve final concentrations of 2, 6, 18, 40, or 100 mM. The reaction was done in replicates and absorbance was measured at 405 nm and 595 nm using a microplate reader operating in kinetic mode at 0, 5, 10, 15, 20, 25, and 30 min intervals.

The assays were prepared at room temperature besides when subjected to heating. The obtained absorbance data showed good difference in presence and absence of DNA conditions at 18 mM concentration. Further studies were done with 20 mM concentration, and it was found to be the most optimal concentration for DNA detection.

**Optimization of methylene blue concentration:** The optimized pH (pH 7.0 and 7.6), salt (50 mM NaCl), experimental condition (with heating) and optimized ferricyanide (20 mM) were used to optimize the concentration of methylene blue for enhanced difference between presence and absence of DNA. Several varied concentrations of methylene blue (10 µM, 20 µM, 40 µM, 80 µM and 100 µM) were being analyzed in the assay with Tris-HCl buffer (pH 7.0 and pH 7.6) for two vital conditions: DNA present ((+)DNA) and DNA absent ((-)DNA). For each different methylene blue concentration, 70 µL (final) of reaction assay was being prepared in replicates, containing the Tris-HCl buffer (final 20 mM), NaCl (final 50 mM), methylene blue and +/-DNA (25 ng/µL). The reaction assay (before ferricyanide addition) was heated at 95°C for 5 mins using a heat block. 63 µL of all the assays were being then transferred to the 96-well plate in three or more replicates, and instantaneously 7 µL of potassium ferricyanide [$K_3Fe(CN)_6$] was added and carefully mixed. Absorbance was being measured at 405 nm and 595 nm using the microplate reader. The kinetic loop was set to record absorbances at multitudinous time-series: 0, 5, 10, 15, 20, 25 and 30 minutes; whilst there was no removal of the plate from the reader during the entirety of absorbance measurement. The assays were being prepared at room temperature except for when subjected to heating. The resultant absorbance data elucidated a pronounced difference in presence and absence of DNA conditions at 80 µM concentration. The difference was indicative of the presence/ absence of DNA and how methylene blue concentration effects the sensitivity of the detection.

**Optimization of ascorbic acid concentration:** The optimized pH (pH 7.0 and 7.6), salt (50 mM NaCl), experimental condition (with heating), optimized ferricyanide (20 mM) and optimized methylene blue (80 µM) were used to optimize the concentration of ascorbic acid (adjuvant) for signal amplification and thereby enhance difference between presence and absence of DNA. Varying concentrations of ascorbic acid (1 mM, 10 mM and 50 mM) were being assessed in the assay with Tris-HCl buffer (pH 7.0 and pH 7.6) for two crucial conditions: DNA present ((+)DNA) and DNA absent ((-)DNA). For each different ascorbic acid concentration, 70 µL (final) of reaction assay was formulated in replicates, containing the Tris-HCl buffer (final 20 mM), NaCl (final 50 mM), methylene blue (80 µM) and +/-DNA (25 ng/µL). The reaction assay before addition of ferricyanide addition was heated at 95°C for 5 mins using a heat block. 63 µL of all the assays were being then transferred to the 96-well plate in three or more replicates, and promptly 7 µL of potassium ferricyanide [$K_3Fe(CN)_6$] was added and carefully mixed. The absorbances were being measured at 405 nm and 595 nm using the microplate reader with the kinetic loop was set to record absorbances at multiple time points: 0, 5, 10, 15, 20, 25 and 30 minutes; whilst there was no removal of the plate from the reader during the entirety of absorbance measurement. The assays were being prepared at ambient room temperature except for when subjected to heating. The obtained absorbance readings showed difference in presence and absence of DNA conditions. This difference reflected the presence/ absence of DNA and signal amplification effects of adjuvant on the detection.

**Nucleic Acid Amplification Tests (NAATs):** The optimized pH (pH 7.0 and 7.6), salt (50 mM NaCl), experimental condition (with heating), optimized ferricyanide (20 mM) and optimized methylene blue (80 µM) and optimized adjuvant concentration (10mM) were used to optimize for compatibility with Nucleic Acid Amplification Tests (NAATs) and hence amplify for enhanced difference between presence and absence of DNA. We also analyzed the effect of adjuvant on Polymerase Chain Reaction (PCR) amplification technique, in order to deduce if adjuvant is also compatible with the NAAT techniques. We conducted the experiment with our novel reaction assay in presence and absence of DNA (+/- 25 ng/µL) for a course of 30 minutes for both without the adjuvant and with the adjuvant.

The reaction was performed in a 30 µL volume, where the template was the 1 µL *E. coli gDNA.* The elution was added with 2 ×Taq mastermix (7.5 µL), forward (5′-AGAGTTTGATCATGGCTCAG-3′) and reverse primers(5-′GGTTACCTTGTTACGACTT-3′) (final concentration was 0.3 µM), and molecular-grade water. The primers were against the E. coli 16S rRNA gene (Accession no: NZ_CP080399.1). For amplification, the cycle was being set at 95 ◦C for 180 s, then at 45 cycles of 95 ◦C for 10 s, 52 ◦C for 10 s, and at 72 ◦C for 30 s, wherein the last step included the amplification analysis.

Glucometer readings of each of the assay conditions were being recorded for 0 , 10 and 30 minutes; whilst the readings were being taken instantly after the NAAT reaction got completed. The assays were being prepared at ambient room temperature except for when subjected to pre-heating for DNA uncoiling and for PCR thermocycling. The obtained glucometer readings showed amplified differential readings in presence and absence of DNA conditions. It also showed that presence of adjuvant amplified the signal difference between with and without DNA assays and that the adjuvant is compatible with NAATs and doesn't degrade due to high temperatures at thermocycling. This difference reflected the presence/absence of DNA and signal amplification effects of adjuvant and PCR on the detection.

**Limit of detection of Polymerase Chain Reaction (PCR):** The optimized pH (pH 7.0 and 7.6), salt (50 mM NaCl), experimental condition (with heating), optimized ferricyanide (20 mM) and optimized methylene blue (80 µM) and optimized adjuvant concentration (10mM) were used to analyze the limit of detection via the PCR nucleic acid amplification technique. LOD and reproducibility assessment of PCR-based detection of nucleic acid was carried out for understanding the glucometer response in range of $10^0$ – $10^8$ copies/reaction of *E coli* gDNA for measurements taken after 0 min, 10 min and 30 min post ferricyanide addition. The assays were being prepared at ambient room temperature except for when subjected to pre-heating for DNA uncoiling and for PCR thermocycling. The obtained glucometer readings showed amplified differential readings in presence and absence of DNA conditions on a range of DNA copies. It also showed that limit of detection is broad and not just limited to single type of copy number. This difference was indicative of the presence/absence of DNA and signal amplification effects of PCR on the detection and its maximum and minimum limit of detection.

**Procedure for Rolling Circle Amplification (RCA) and detection via glucometer:** The optimized pH (pH 7.0 and 7.6), salt (50 mM NaCl), experimental condition (with heating), optimized ferricyanide (20 mM) and optimized methylene blue (80 μM) and optimized adjuvant concentration (10mM) were used to analyze the compatibility with endpoint RCA nucleic acid amplification technique.

**2.3.1 5′-Phosphorylation**: Precursor ssDNA oligonucleotides were diluted with DEPC treated water to 8 μM. The oligonucleotide solution was snap-cooled (5 minutes heating at 95°C immediately followed by 5 minutes incubation on ice) to linearize the DNA for improving the phosphorylation efficiency. The 5′-phosphorylation was carried on the oligonucleotides (final concentration 4 μM) in presence of T4 PNK (0.375 Units μL-1), ATP (1 mM), ligase buffer (50 mM Tris-HCl pH 7.5, 10 mM MgCl2, 1 mM ATP, 10 mM DTT), DTT (5 mM), and spermidine (1.7 mM). The addition of all the reagents was performed on ice. The solution was incubated at 37°C for 3 hours followed by annealing by slow (1 h) cooling from 95°C to 4°C (this step also inactivated the T4 PNK). For sticky-end substrates, precursor oligonucleotides (**3a**/**3b** or **4a**/**4b**) were separately 5′-phosphorylated, mixed 1:1 (v/v, to a final concentration 2 μM), and then subjected to annealing as described above. For splint-padlock ligation, phosphorylation was carried out exactly as described above followed by T4 PNK inactivation (75°C for 20 min). Following the mixing of splint (final concentration 3.84 μM) with 5′-phosphorylated padlock (final concentration 3.2 μM), the solution was subjected to annealing as described above.

**Ligation:** Circularization of the annealed 5′-phosphorylated DNA (concentration 3.2 μM for self-annealing precursor, 1.6 μM for sticky-end precursors and 3.2 μM for the padlock) was carried out in presence of T4 DNA ligase (8 Cohesive End Units μL-1), ATP (1 mM), and T4 DNA ligase buffer (50 mM Tris-HCl pH 7.5, 10 mM MgCl2, 1 mM ATP, 10 mM DTT). The addition of all the reagents was done on ice. The ligation mixture was then incubated at 16°C for 16 hours followed by enzyme inactivation at 75°C for 20 min.

**Exonucleases treatment:** The ligated sample was snap-cooled as described in Section 2.3.1. The exonuclease digestion was carried out in 25 μL reaction volume as described below with final oligonucleotide concentrations 1.5 μM for self-annealing ligation, 0.75 μM for sticky-end ligation, and 1.5 μM for self-annealing ligation. For exclusive exonuclease I or III digestion, the reaction was carried out with the appropriate exonuclease enzyme (2.4 units $\mu L^{-1}$ for exonuclease I or 12 units $\mu L^{-1}$ for exonuclease III) in exonuclease I buffer (67 mM Glycine-KOH pH 9.5, 6.7 mM MgCl2, 10 mM β-mercaptoethanol) or exonuclease III buffer (10 mM Bis-Tris-Propane-HCl pH 7, 10 mM MgCl2, 1 mM DTT). For double digestion, both exonuclease I (2.4 units μL-1) and III (12 units μL-1) were used in exonuclease III buffer (10 mM Bis-Tris-Propane-HCl pH 7, 10 mM MgCl2, 1 mM DTT). All the reactions were supplemented with DTT (1 mM) to ensure complete digestion. The reagents were added while keeping the vials on ice. The solutions were incubated at 37°C for 4 hours, followed by enzyme inactivation at 85°C for 20 minutes and then stepwise annealing from 85°C to 4°C to generate the desired secondary structure.

**Endpoint RCA**

Before initiation endpoint RCA reactions, the oligonucleotide substrates were incubated at 30°C for 30 min in 50 mM Tris-HCl (pH 8) and 50 mM NaCl. The endpoint RCA reactions was carried out in 30 μL volume in presence of circular DNA (0.02 µM), (optional) Primer and/or endpoint RCA primer (0.02 µM), dNTPs (1.0 mM), BSA (0.2 μg uL-1), ϕ29 buffer (50 mM Tris-HCl pH 7.5, 10 mM MgCl2, 10 mM (NH4)2SO4, 4 mM DTT), Methylene blue (80 μM), and ϕ29 DNA polymerase (0.2 units μL-1). The reactions were incubated for 2h at 30°C and fluorescence intensities were recorded in 2 min intervals.

**Glucometer Measurement for RCA**

After the completion of RCA, ferricyanide (20 mM) was being instantly added to the assay and then glucometer readings were being taken for the time-period course of 0 minute, 10 minutes and 30 minutes for both presence and absence of DNA assays. The assays were being prepared at ambient room temperature except for when subjected to pre-heating for DNA uncoiling and for endpoint RCA temperature conditions. In addition to that the glucometer readings were also being taken at the ambient room temperature in a dark room facility. The recorded glucometer readings indicated amplified and enhanced differential readings in presence and absence of DNA conditions. This experiment showed that our novel assay pipeline is compatible with Rolling Circle Amplification (RCA) and also showed the signal amplification effects of RCA as evident in the enhanced glucometer signal difference between DNA-present and DNA-absent conditions.

**Procedure for Loop-mediated Isothermal Amplification (LAMP) and detection via glucometer:** The optimized pH (pH 7.0 and 7.6), salt (50 mM NaCl), experimental condition (with heating), optimized ferricyanide (20 mM) and optimized methylene blue (80 µM) and optimized adjuvant concentration (10mM) were used to analyze the compatibility with endpoint LAMP nucleic acid amplification technique.

### 2.2. LAMP reaction and primer optimization using real-time fluorescence readout

A endpoint LAMP experiment was performed on $10^3$ copies of ORF1ab containing plasmid with three sets of primers. The final LAMP reaction (30 µl) contained the three primer pairs in the following final concentrations: 0.2 µM outer primers, 1.6 µM forward and backward inner primers [27,28]. For primer set 2, 0.4 µM outer primers, 0.332 µM forward and backward inner primers were utilized in the final concentration [29]. The reaction mixture also contained 2.0 µL of 10 × Bst 2.0 DNA polymerase reaction buffer [1 × containing 20 mM Tris-HCl, 50 mM KCl, 10 mM $(NH_4)_2SO_4$,2 mM $MgSO_4$, 0.1% Tween-20, pH 8.8], 1.4 mM dNTPs, Methylene blue (final concentration 80 µM), 0.5 µL of an 8 U µl concentration of Bst 2.0 DNA polymerase, 6 mM $MgSO_4$ and 1 µl template (alternatively, 2 µL magnetic bead for magnetocapture assays). For endpoint LAMP, 7 U (0.25 µL) of reverse transcriptase RTx (NEB) was additionally added to the above. The LAMP reaction was set at the following settings for each cycle with a fluorescence monitoring step, 65 °C for 1 min for primer set 1, 64 °C for 1 min for primer set 2, 60 °C for 1 min for primer set 3 followed by thermal melting analysis step. The cycles were repeated 60 times (unless otherwise stated) in a CFX Maestro or CFX Connect real-time PCR (rt-PCR) machine (BioRad).

### 2.3. LAMP assays

For endpoint LAMP using the pure nucleic acid template, the assay was performed on $10^1$–$10^4$ copies of nucleic acid (*rdp gene of COVID-19*)/25 μL of reaction. For magnetocapture followed LAMP assays, 2 μL magnetic beads containing immobilized target nucleic acid were added to 25 μL electrochemical LAMP reaction having a composition as described below. A 25 μL LAMP reaction comprised of 2.5 μL of 10 × Bst 2.0 DNA polymerase reaction buffer [1 × containing 20 mM Tris-HCl, 50 mM KCl, 10 mM $(NH_4)_2SO_4$, 2 mM $MgSO_4$, 0.1% Tween-20, pH 8.8], 1.4 mM dNTPs, 0.4 μM outer primers, 0.332 μM forward and backward inner primer, 1 μM forward loop primers, 0.4 μM back loop primers (primer set 2), 0.5 μL Bst 2.0 polymerase, and 50 μM methylene blue. During magnetocapture followed by eLAMP or eRT-LAMP, 1000 copies of non-magnetocaptured plasmid DNA or RNA was used as the positive control. For LAMP, 7 U of reverse transcriptase (RTx) was also used for 25 μL of reaction. The assays were set at the following settings for each cycle, 64 °C for 1 min for 60 cycles followed by heat inactivation at 80 °C for 20 min in a thermal cycler (Eppendorf). The resultant LAMP amplicons were analyzed via glucometer readings.

**Glucometer Measurement for LAMP**

After the completion of LAMP, ferricyanide (20 mM) was being instantly added to the assay and then glucometer readings were being taken for the time-period course of 0 minute, 10 minutes and 30 minutes for both presence and absence of DNA assays. The assays were being prepared at ambient room temperature except for when subjected to pre-heating for DNA uncoiling and for endpoint LAMP temperature conditions. In addition to that the glucometer readings were also being taken at the ambient room temperature in a dark room facility. The recorded glucometer readings showed evident amplified and enhanced differential readings in DNA-present and DNA-absent conditions. This experiment indicated that our novel assay pipeline is compatible with Loop-mediated Isothermal Amplification (LAMP) and also showed the signal amplification effects of LAMP as shown in the enhanced glucometer signal difference between DNA(+) and DNA(-) conditions.

**Glucometer measurement:** The mixture of the reagents was same as mentioned above. Majority of the experiments were being carried out using the “Dr. Morepen GLuco One” brand, while the investigation with an alternative brand was performed with “Apollo pharmacy blood glucose monitoring” brand. At previously specified time point, 1 μL of the reaction was aliquoted onto a hydrophobic parafilm strip. The glucometer measurement was then carried out as instructed in the manufacturer's manual using the 1 μL droplet. In case of the Dr. Morepen GLuco One or Apollo pharmacy blood glucose monitoring, it involved first inserting the strip electrode into the glucometer device, and then waiting for the blood droplet image to appear on screen (within 2-3 seconds), and then bringing the designated place for blood droplet insertion in contact with the reaction droplet and then readout is noted.

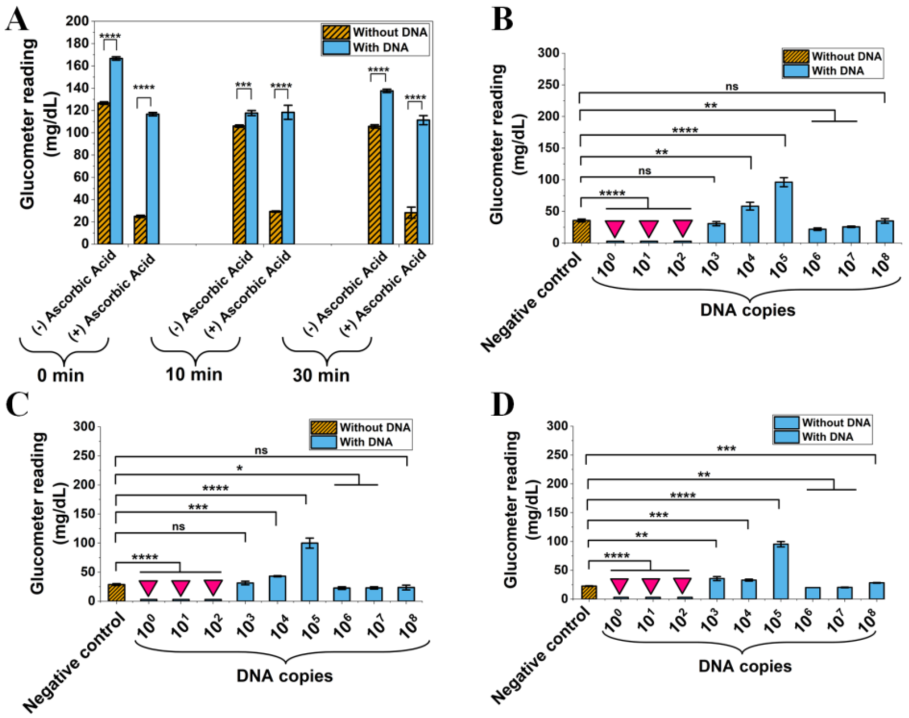


**Figure S1.** Glucometer-based assessment of PCR (with and without DNA template) and LOD for detecting *E coli* gDNA using MB (80 μM), potassium ferricyanide (20 mM), and ascorbic acid (10 mM). A, Time point (0, 10, 30 min) analysis of glucometer response for PCR reaction on $10^5$ copies/reaction of *E coli* gDNA template as a function of ascorbic acid presence. Measurements were taken after 0 min, 10 min, or 30 min after ferricyanide addition. B – D, Glucometer response for detecting $10^0$ – $10^8$ copies/reaction of *E coli* gDNA for measurements taken after 0 min (panel B), 10 min (panel C), and 30 min (panel D) of ferricyanide addition. Error bars represent standard deviation (n = 4). Statistical analysis performed using Student's t-test. $P < 0.05$ (*), $P < 0.01$ (**), $P < 0.001$ (***), $P < 0.0001$ (****).
